# The SARS-CoV-2 multibasic cleavage site facilitates early serine protease-mediated entry into organoid-derived human airway cells

**DOI:** 10.1101/2020.09.07.286120

**Authors:** Anna Z. Mykytyn, Tim I. Breugem, Samra Riesebosch, Debby Schipper, Petra B. van den Doel, Robbert J. Rottier, Mart M. Lamers, Bart L. Haagmans

**Affiliations:** Viroscience Department, Erasmus University Medical Center, Rotterdam, the Netherlands; Department of Pediatric Surgery, Erasmus University Medical Center - Sophia Children’s Hospital, Rotterdam, the Netherlands

**Keywords:** COVID-19, SARS-CoV-2, SARS-CoV, multibasic cleavage site, serine proteases, airway organoids, furin site

## Abstract

After the SARS-CoV outbreak in 2003, a second zoonotic coronavirus named SARS-CoV-2, emerged late 2019 in China and rapidly caused the COVID-19 pandemic leading to a public health crisis of an unprecedented scale. Despite the fact that SARS-CoV-2 uses the same receptor as SARS-CoV, transmission and pathogenesis of both viruses seem to be quite distinct. A remarkable feature of the SARS-CoV-2 spike is the presence of a multibasic cleavage site, which is absent in the SARS-CoV spike. The viral spike protein not only attaches to the entry receptor, but also mediates fusion after cleavage by host proteases. Here, we report that the SARS-CoV-2 spike multibasic cleavage site increases infectivity on differentiated organoid-derived human airway cells. Compared with SARS-CoV, SARS-CoV-2 entered faster into the lung cell line Calu-3, and more frequently formed syncytial cells in differentiated organoid-derived human airway cells. Moreover, the multibasic cleavage site increased entry speed and plasma membrane serine protease usage relative to endosomal entry using cathepsins. Blocking serine protease activity using the clinically approved drug camostat mesylate effectively inhibited SARS-CoV-2 entry and replication in differentiated organoid-derived human airway cells. Our findings provide novel information on how SARS-CoV-2 enters relevant airway cells and highlight serine proteases as an attractive antiviral target.

**Significance Statement:** Highly pathogenic coronaviruses have spilled from animals to humans three times in the past two decades. Late 2019, SARS-CoV-2 emerged in China and was declared a pandemic by March 2020. The other two highly pathogenic coronaviruses, SARS-CoV and MERS-CoV, emerged in 2002 and 2012, respectively, but did not attain sustained human-to-human transmission. Given the high diversity of coronaviruses in animals, urbanization and increased air travel, future coronavirus pandemics are likely to occur intermittently. Identifying which factors determine pandemic potential and pathogenicity are therefore of key importance to global health. Additionally, there is an urgent need to rapidly translate fundamental knowledge to the clinic, a process that is expedited through the use of relevant cell culture systems.

## Introduction

The ongoing coronavirus disease (COVID-19) pandemic is caused by the severe acute respiratory syndrome coronavirus 2 (SARS-CoV-2), which emerged in central China late 2019 (1). Within months this virus spread globally and as of August 26, 2020, over 24 million cases have been reported, including over 800,000 deaths. Halting SARS-CoV-2 spread has shown to be highly complex, putting great strain on health systems globally. SARS-CoV-2 is the third zoonotic coronavirus to emerge from animal reservoirs within the past two decades, after SARS-CoV and Middle East respiratory syndrome coronavirus (MERS-CoV), in 2002 and 2012, respectively (2–5). In contrast to SARS-CoV-2, SARS-CoV and MERS-CoV have not attained sustained human-to-human transmission. These coronaviruses belong to the *Betacoronavirus* genus (family *Coronaviridae*, subfamily *Orthocoronavirinae*), which is thought to ultimately originate from bats, but can spread to humans via intermediate hosts (6–8).

Currently it is largely unknown what factors determine coronavirus transmission to and between humans, but one important determinant may be the coronavirus spike (S) protein, which is the main glycoprotein incorporated into the viral envelope. Enveloped viruses, including coronaviruses, deposit their genomes into host cells by coalescing their membranes with the cell. This function is executed by S protein trimers, which fuse viral and cellular membranes after binding to the entry receptor (9). In addition, coronaviruses can spread from cell to cell when coronavirus S proteins traffic to the plasma membrane of infected cells and fuse with neighboring cells, generating multinucleated giant cells (syncytia). Coronavirus S proteins are synthesized in infected cells in a stable and fusion-incompetent form and are activated through cleavage by host proteases. Proteolysis controls the timely release of the S protein’s stored energy required to fuse membranes, which allows virions to be stable in the environment yet fusogenic after contacting entry receptors on host cell membranes.

Cleavage is essential for coronavirus infectivity and can occur in the secretory pathway of infected cells or during viral entry into target cells (9, 10). Several groups of host proteases, including type II transmembrane serine proteases (hereafter referred to as serine proteases), proprotein convertases and cathepsins, can cleave the S protein. Specific sites in the S protein regulate protease usage and therefore play an important role in determining cell tropism. Similarly, tropism can be determined by the availability of proteases that can activate the S protein (10–14). The S protein consists of two domains, the receptor binding (S1) domain and the fusion (S2) domain. These domains are separated by the S1/S2 cleavage site, which in some coronaviruses such as SARS-CoV-2, forms an exposed loop that harbors multiple arginine residues and is therefore referred to as a multibasic cleavage site (15, 16). Cleavage of this site can occur in secretory systems of infected cells by proprotein convertases, including furin. S1/S2 cleavage does not directly trigger fusion but may facilitate or regulate further cleavage (17). A second proteolysis step takes place at a more C-terminal site within the S2 domain, notably the S2’ site. S2’ cleavage is thought to occur after the virus has been released from producing cells and is bound to host cell receptors on receiving cells. The S2’ site is processed by serine proteases on the plasma membrane or by cathepsins in the endosome. Whereas S2’ cleavage appears to be crucial for coronavirus infectivity, not all coronaviruses contain a multibasic S1/S2 site and little is known of its function (9). Until recently, all viruses within the clade of SARS-related viruses, including SARS-CoV, were found to lack a multibasic S1/S2 cleavage site. However, SARS-CoV-2 contains a PRRA insertion into the S protein, precisely N-terminally from a conserved arginine, creating a multibasic RRAR cleavage motif. Exchanging the SARS-CoV-2 S multibasic cleavage site for the SARS-CoV monobasic site was recently shown to decrease fusogenicity on a monkey kidney cell line (VeroE6) and infectivity in a human lung adenocarcinoma cell line (Calu-3) (18, 19). However, cancer cells often poorly represent untransformed cells and thus the question remains whether the multibasic cleavage site would affect infectivity on relevant lung cells. Another study showed that entry of SARS-CoV-2 pseudoparticles into both Calu-3 cells and primary airway cultures could be blocked using a clinically approved serine protease inhibitor (camostat mesylate), but no effects on authentic virus entry and replication were shown (20). Here we investigated if the SARS-CoV-2 multibasic cleavage site affects entry into relevant human lung cells (i), if the multibasic cleavage site can alter protease usage during entry (ii), and if authentic SARS-CoV-2 entry and replication can be inhibited using camostat mesylate (iii).

## Results

### Entry into lung adenocarcinoma and differentiated organoid-derived human airway cells is facilitated by the SARS-CoV-2 S multibasic cleavage site

Recently, Hoffmann and colleagues (2020) showed that the SARS-CoV-2 multibasic cleavage motif increases entry into Calu-3 cells by exchanged this motif and several N-terminally flanking amino acids with the monobasic S1/S2 site found in SARS-CoV or in a related bat virus RaTG13 (18). Building on these observations, we generated several SARS-CoV-2 S protein mutants and used these to generate vesicular stomatitis virus- (VSV) based pseudoparticle stocks (PPs) expressing a green fluorescent protein (GFP). Instead of exchanging cleavage sites, we mutated the minimal RXXR multibasic cleavage motif by deleting the PRRA insertion (Del-PRRA), changing the last arginine to an alanine (R685A), or to a histidine (R685H) in order to preserve the positive charge at this site (Fig. 1A). Immunoblotting revealed that wild type and mutant PPs were produced at similar levels (Fig. 1B). S1/S2 cleavage was observed for the wild type SARS-CoV-2 PPs and abrogated by the PRRA deletion and the R685A and R685H substitutions, which is in agreement with studies showing that the SARS-CoV-2 S is cleaved by proprotein convertases, possibly furin (18, 19). Next, we assessed the infectivity of these viruses and found that the SARS-2-Del-PRRA and SARS-2-R685A mutants were 5-10 fold more infectious on VeroE6 cells (Fig. 1C). In contrast, on the lung adenocarcinoma cell line Calu-3 the SARS-2-Del-PRRA, SARS-2-R685A, and SARS-2-R685H PPs were approximately 5-10 fold less infectious compared with the wild type PPs (Fig. 1D). These data show that the PRRA deletion and single point mutations could functionally destroy the multibasic cleavage site and suggest that this site enhances lung cell entry. Next, we assessed the effect of the multibasic cleavage site in a relevant cell culture system, using airway organoids (21) that were dissociated, seeded onto collagen coated Transwell inserts and differentiated for 10-11 weeks at air-liquid interface in Pneumacult ALI medium (Stemcell). After differentiation, cultures contained ciliated cells, club cells and goblet cells (Fig. S1A-C). Moreover, they expressed the SARS-CoV-2 entry receptor angiotensin converting enzyme 2 (ACE2) and transmembrane protease serine 2 (TMPRSS2), a serine protease previously shown to mediate SARS-CoV-2 entry when overexpressed (Fig. S1D-E) (20). For infection experiments, the differentiated pseudostratified epithelial layer was dissociated into small clumps, infected in suspension and then re-plated into basement membrane extract (BME), in which it formed spheroids. SARS-CoV-2 PPs successfully infected airway spheroids, as observed by fluorescent microscopy (Fig. 1E). SARS-CoV-2 PPs were approximately 2 times more infectious on these cells compared with the SARS-2-Del-PRRA and SARS-2-R685A mutants, and 8 times more infectious than the SARS-2-R685H mutant (Fig. 1F), demonstrating that the SARS-CoV-2 multibasic cleavage site facilitates entry into relevant human airway cells.

**Figure 1.**
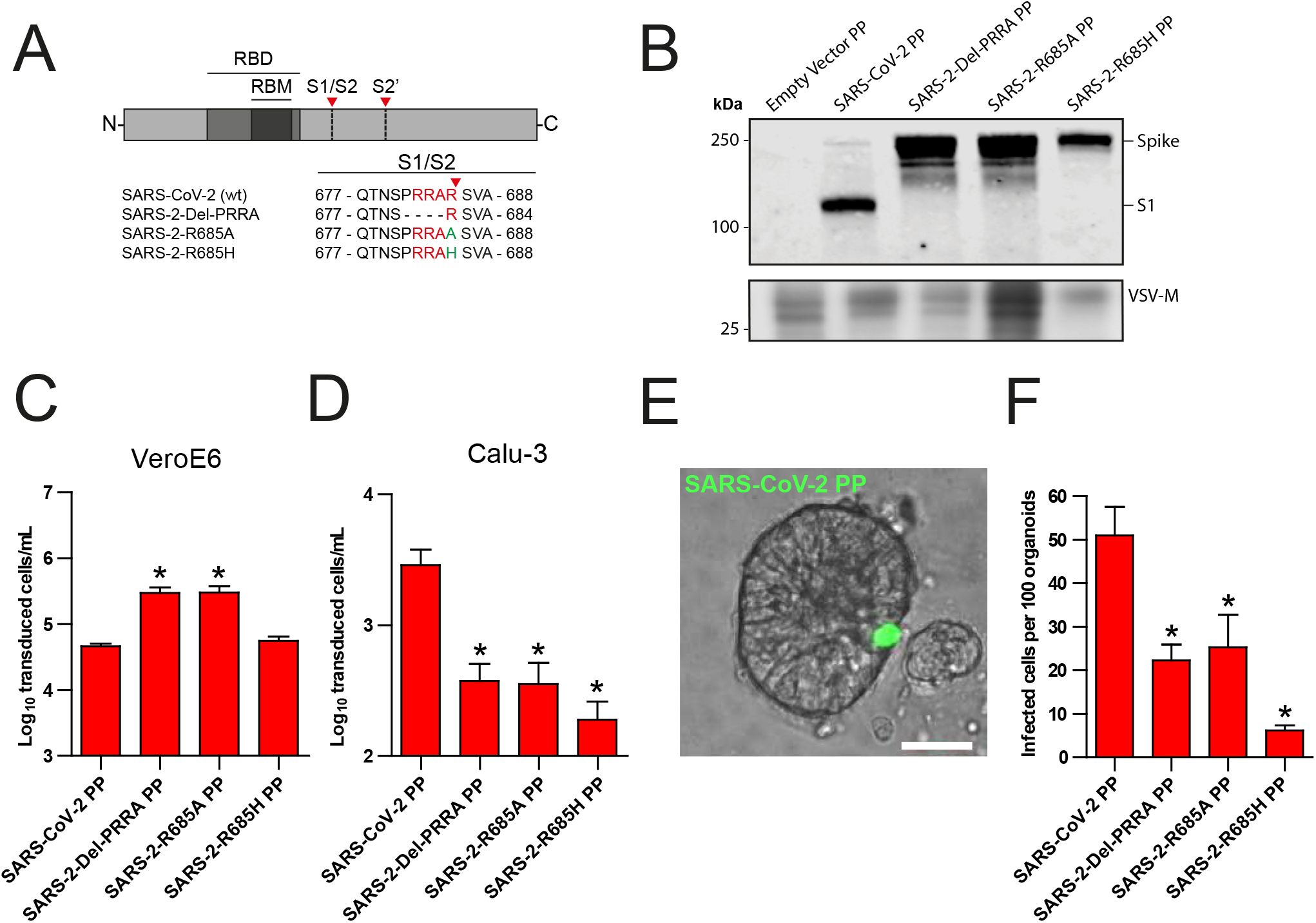
The SARS-CoV-2 S multibasic cleavage site mediates entry into organoid-derived human airway cells. (*A*) Schematic overview of SARS-CoV-2 S protein mutants. Multibasic cleavage site residues are indicated in red; amino acid substitutions are indicated in green. Red arrows indicate cleavage sites. RBD = receptor binding domain, RBM = receptor binding motif. The SARS-CoV-2 S multibasic cleavage site was mutated to either remove the PRRA motif (SARS-2-Del-PRRA) or to substitute the R685 site (SARS-2-R685A and R685H). (*B*) Comparison of S cleavage of SARS-CoV-2 PPs and the multibasic cleavage site mutants. Western blots were performed against S1 with VSV-M silver stains as a production control. (*C* and *D*) PP infectivity of SARS-CoV-2 S and multibasic cleavage site mutants on VeroE6 (*C*) and Calu-3 (*D*) cells. (*E*) Differentiated airway spheroid cultures were infected with concentrated SARS-CoV-2 PPs containing a GFP reporter, indicated in green. Scale bar indicates 20 μm. (*F*) SARS-CoV-2 PP and multibasic cleavage site mutant infectivity on differentiated bronchiolar airway spheroid cultures. One-way ANOVA was performed for statistical analysis comparing all groups with SARS-CoV-2 PPs. * P<0.05. Error bars indicate SEM. PP = pseudoparticles. Experiments were performed in triplicate (*C* and *D*, *F*). Representative experiments from at least two independent experiments are shown.

### SARS-CoV-2 enters Calu-3 cells faster than SARS-CoV and entry speed is increased by the multibasic cleavage site

As SARS-CoV lacks the multibasic cleavage site, we compared its infectivity to SARS-CoV-2 and found that both PPs readily infected Calu-3 cells, indicating that the SARS-CoV S has adaptations other than the multibasic cleavage site to facilitate airway cell infection (Fig. 2A). Likewise, inserting the PRRA motif into SARS-CoV S and thereby generating a multibasic cleavage site did not increase PP infectivity on Calu-3 cells (Fig. S2). To investigate this further, we compared the entry route taken by these viruses in Calu-3 cells. For this purpose, we used inhibitors of two major coronavirus entry pathways (9). Serine proteases are known to mediate early coronavirus entry on the plasma membrane or in the early endosome, whereas cathepsins facilitate entry in late, acidified endosomes. Concentration ranges of either a serine protease inhibitor (camostat mesylate; hereafter referred to as camostat) or a cathepsin inhibitor (E64D) were used to assess the entry route into Calu-3 cells. Entry of SARS-CoV-2 PPs was not inhibited by E64D, but could be inhibited by camostat, indicating that SARS-CoV-2 exclusively uses serine proteases for entry into these cells (Fig. 2B-C). For SARS-CoV PPs, entry was inhibited slightly by E64D (~10%), but camostat had a far stronger effect (~90%), indicating that SARS-CoV mainly uses serine proteases to enter Calu-3 cells but that a small fraction of virions enter via cathepsins (Fig. 2B-C). Previously, Calu-3 cells have been suggested to have low levels of cathepsin activity (16). The observation that some SARS-CoV PPs use cathepsins suggests that this virus less efficiently uses the surface serine proteases encountered early during entry, resulting in particles accumulating in the endosome, where they are cleaved by cathepsins. To test this, we measured the serine protease-mediated entry rate of SARS-CoV-2 and SARS-CoV by blocking entry on Calu-3 cells at different time points post infection using camostat. Cells were pretreated with E64D to prevent any cathepsin-mediated entry. Using both PPs and authentic virus (Fig. 2D-E) we observed that SARS-CoV-2 entered faster than SARS-CoV via serine proteases. Next, we assessed whether the presence of a multibasic cleavage site could increase the serine protease-mediated entry rate into Calu-3 cells. For this purpose, we used SARS-CoV S PPs containing the PRRA insertion (SARS-PRRA) (Fig. 2F). Immunoblotting revealed that, in contrast to SARS-CoV PPs, SARS-PRRA PPs were partially cleaved (Fig. 2G). Whereas wild type SARS-CoV PPs used cathepsins, SARS-PRRA PPs did not (Fig. 2H-I). The serine protease-mediated entry rate of SARS-PRRA PPs on Calu-3 cells was higher compared with SARS-CoV PPs (Fig. 2J), and it was lower for SARS-2-Del-PRRA PPs compared with SARS-CoV-2 PPs (Fig. 2K). These findings show that the SARS-CoV-2 multibasic cleavage site facilitates serine protease-mediated entry on Calu-3 cells.

**Figure 2.**
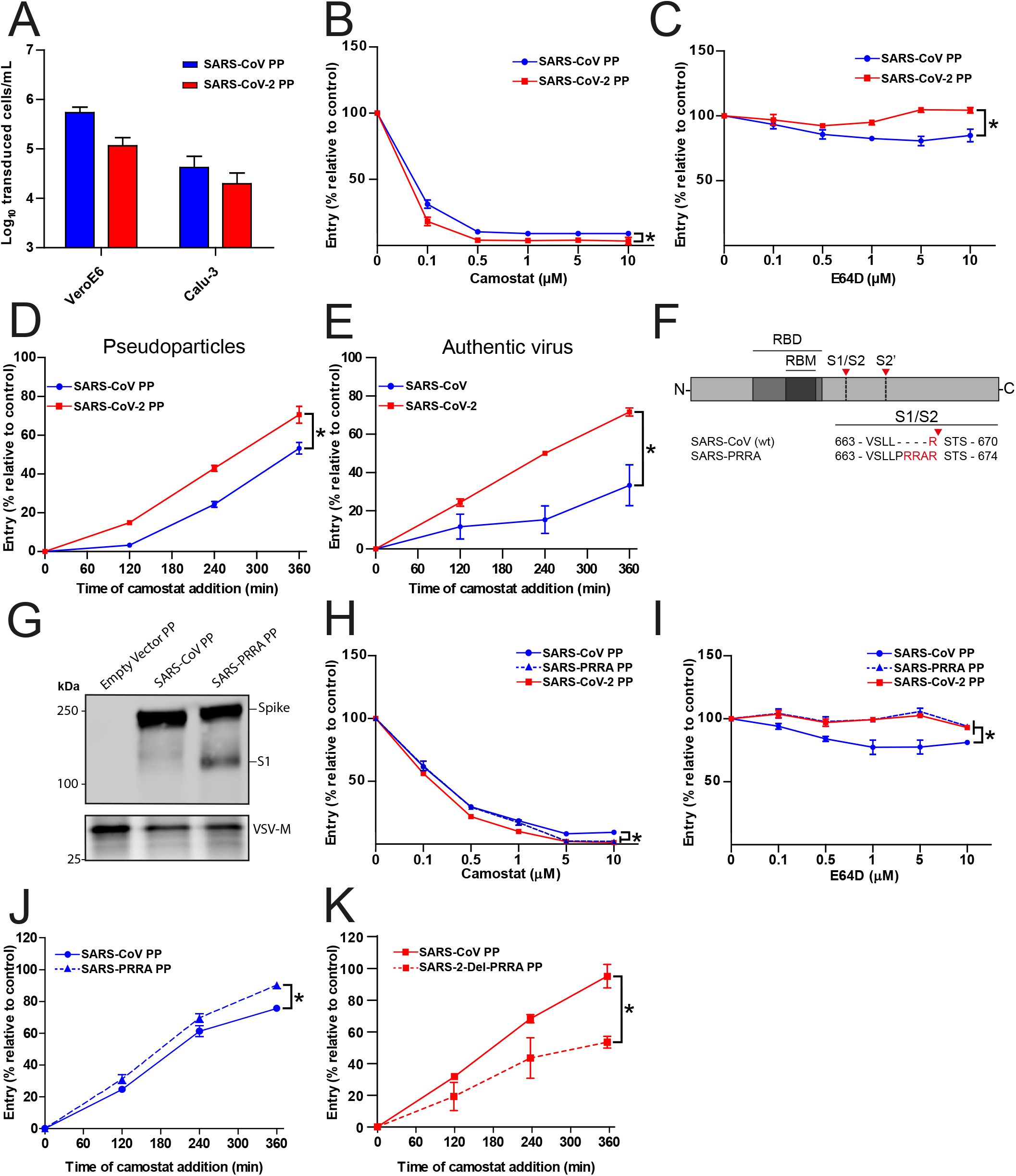
SARS-CoV-2 enters faster on Calu-3 cells than SARS-CoV and entry speed is increased by the multibasic cleavage site. (*A*) SARS-CoV PP and SARS-CoV-2 PP infectivity on VeroE6 and Calu-3 cells. (*B* and *C*) SARS-CoV PP and SARS-CoV-2 PP entry route on Calu-3 cells. Cells were pretreated with a concentration range of camostat (*B*) or E64D (*C*) to inhibit serine proteases and cathepsins, respectively. T-test was performed for statistical analysis at the highest concentration. * P<0.05. (*D* and *E*) SARS-CoV PP, SARS-CoV-2 PP (*D*) and authentic virus (*E*) entry speed on Calu-3 cells. T-test was performed for statistical analysis at the latest time point. * P<0.05. (*F*) Schematic overview of SARS-CoV S protein mutants. Multibasic cleavage site residues are indicated in red. The SARS-CoV-2 PRRA motif was inserted into SARS-CoV PPs (SARS-PRRA). (*G*) Comparison of S1 cleavage of SARS-CoV PP and the multibasic cleavage site mutant. VSV-M silver stains are shown as a production control. (*H* and *I*) SARS-CoV PP, SARS-PRRA PP and SARS-CoV-2 PP entry route on Calu-3 cells. Cells were pretreated with a concentration range of camostat (*H*) or E64D (*I*) to inhibit plasma membrane and endosomal entry respectively. One-way ANOVA was performed for statistical analysis comparing all groups with SARS-CoV PPs at the highest concentration. * P<0.05. (*J* and *K*) Entry speed on Calu-3 cells of SARS-CoV PPs compared with SARS-PRRA PPs (*J*) and SARS-CoV-2 PPs compared with SARS-2-Del-PRRA PPs (*K*). T-test was performed for statistical analysis at the latest time point. * P<0.05. Error bars indicate SEM. PP = pseudoparticles. Experiments were performed in triplicate (*A* to *E, H* to *K*). Representative experiments from at least two independent experiments are shown.

### Cell-cell fusion is facilitated by the SARS-CoV-2 multibasic cleavage site and SARS-CoV-2 is more fusogenic than SARS-CoV on differentiated organoid-derived human airway cells

Next, we used a GFP-complementation cell-cell fusion assay (Fig. S3) to determine whether entry rate was associated with fusogenicity. In this assay, S and GFP-11 co-transfected HEK-293T cells fuse with GFP1-10 expressing Calu-3 cells, resulting in GFP complementation and fluorescence. In HEK-293T cells, multibasic cleavage site containing S proteins were more cleaved than S proteins without this site (Fig. 3A). We observed that SARS-CoV-2 S was more fusogenic than SARS-CoV S on Calu-3 cells (Fig. 3B-C) and in addition, the insertion of the multibasic cleavage site into SARS-CoV S increased fusion, whereas mutations in the SARS-CoV-2 S multibasic cleavage site decreased fusion. To investigate differences in fusogenicity in a relevant cell system, we infected 2D differentiated organoid-derived human airway air-liquid interface cultures with SARS-CoV-2 and SARS-CoV and assessed the formation of syncytial cells at 72 hours post infection using confocal microscopy. Cells were termed syncytial cells when at least two nuclei were present within a single viral antigen positive cell that lacked demarcating tight junctions. SARS-CoV-2 frequently induced syncytia, whereas SARS-CoV-infected cells rarely contained multiple nuclei (Fig. 3D; and E for quantification).

**Figure 3.**
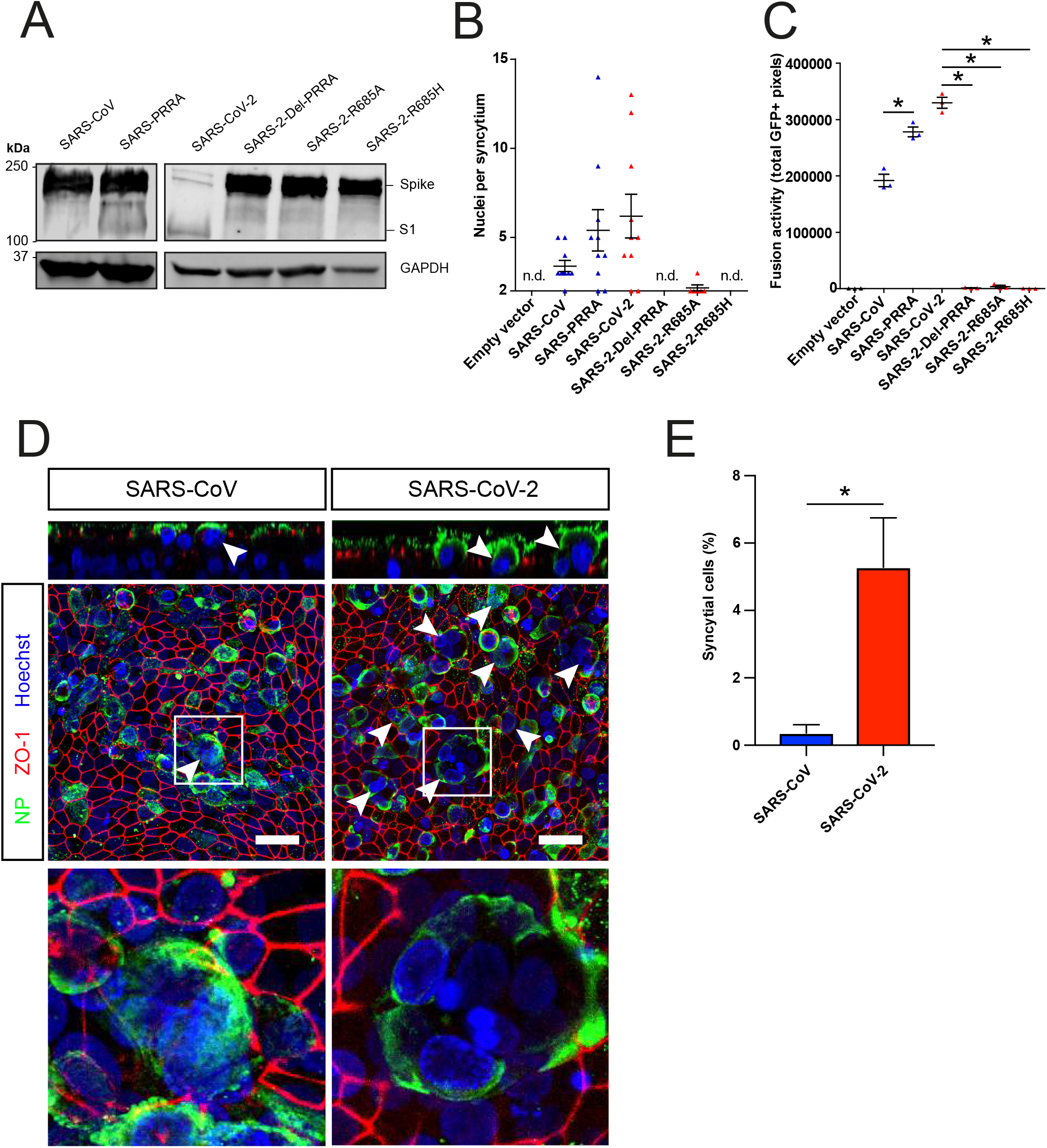
The SARS-CoV-2 multibasic cleavage site facilitates cell-cell fusion and SARS-CoV-2 is more fusogenic than SARS-CoV on differentiated organoid-derived human airway cells. (*A*) Proteolytic cleavage of SARS-CoV-2 S, SARS-CoV S, and S mutants was assessed by overexpression in HEK-293T cells and subsequent western blots for S1. GAPDH was used as a loading control. (*B* and *C*) Fusogenicity of SARS-CoV-2 S, SARS-CoV S, and S mutants was assessed after 18 hours by counting the number of nuclei per syncytium (*B*) and by measuring the sum of all GFP+ pixels per well (*C*). Statistical analysis was performed by one-way ANOVA on SARS-CoV or SARS-CoV-2 S-mediated fusion compared with its respective mutants. * P<0.05 (*C*). (*D*) Differentiated bronchiolar airway cultures were infected at a MOI of 1 with SARS-CoV or SARS-CoV-2. 72 hours post infection they were fixed and stained for nucleoprotein (NP; green) and tight junctions (ZO1; red) to image syncytia. Nuclei were stained with hoechst (blue). Scale bars indicate 20 μm. Arrows indicate syncytial cells. (*E*) Percentage of syncytial cells of total number of infected cells per field of 0.1 square mm. 5 fields were counted. T-test was performed for statistical analysis. * P<0.05. Error bars indicate SEM. Experiments were performed in triplicate (*C*). Representative experiments from at least two independent experiments are shown.

### The SARS-CoV-2 multibasic cleavage site increases serine protease usage and decreases cathepsin usage

The findings above indicate that SARS-CoV-2 S is more fusogenic and mediates faster entry through serine proteases compared with SARS-CoV, indicating that the multibasic cleavage site alters protease usage. To investigate this, cells that contain both serine and cathepsin protease-mediated entry should be used. Therefore, we focused on VeroE6 cells, which have an active cathepsin-mediated cell entry pathway, as on these cells both SARS-CoV-2 PP and SARS-CoV PP entry was blocked by E64D, and not by camostat (Fig. 4A-B). To generate a cell line in which both entry pathways are active, we stably expressed TMPRSS2 in VeroE6 cells. In these cells, SARS-CoV-2 PP entry was inhibited ~95% by camostat, whereas SARS-CoV PPs were only inhibited ~35% (Fig. 4C-D). In accordance, E64D did not block SARS-CoV-2 PP entry, while it decreased SARS-CoV PP entry ~30%. These findings indicate that despite a functional serine protease-mediated entry pathway, a significant part of SARS-CoV PPs still retained cathepsin-mediated entry whereas SARS-CoV-2 PPs only used serine proteases for entry. This phenotype was found to be linked to the multibasic cleavage site as SARS-CoV-2 PPs containing mutations in this site entered less through serine proteases and more through cathepsins (Fig. 4E-F). In accordance, the introduction of the multibasic cleavage site into SARS-CoV PPs increased serine proteases usage, while decreasing cathepsin usage (Fig. 4G-H).

**Figure 4.**
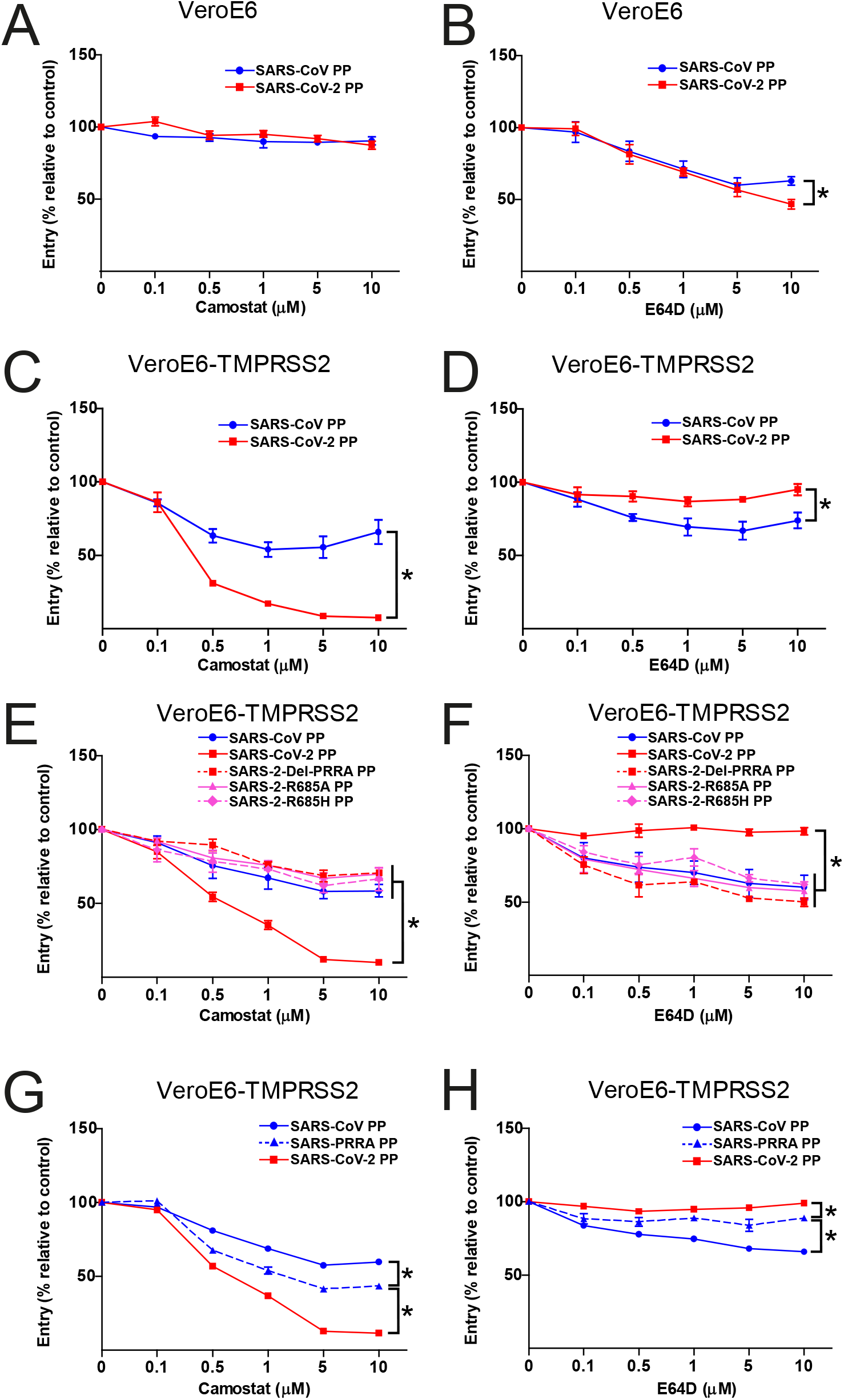
The SARS-CoV-2 multibasic cleavage site increases serine protease usage. (*A* and *B*) SARS-CoV PP and SARS-CoV-2 PP entry route on VeroE6 cells pretreated with a concentration range of camostat (*A*) or E64D (*B*) to inhibit serine proteases and cathepsins, respectively. (*C* and *D*) SARS-CoV PP and SARS-CoV-2 PP entry route on VeroE6-TMPRSS2 cells pretreated with a concentration range of camostat (*C*) or E64D (*D*) to inhibit serine proteases and cathepsins, respectively. T-test was performed for statistical analysis at the highest concentration. * P<0.05. (*E* and *F*) Entry route of SARS-CoV-2 PP and multibasic cleavage site mutants on VeroE6-TMPRSS2 cells pretreated with a concentration range of camostat (*E*) or E64D (*F*) to inhibit serine proteases and cathepsins, respectively. One-way ANOVA was performed for statistical analysis comparing all groups to SARS-CoV-2 PPs at the highest concentration. * P<0.05. (*G* and *H*) Entry route of SARS-CoV PPs and SARS-PRRA PPs on VeroE6-TMPRSS2 cells pretreated with a concentration range of camostat (*G*) and E64D (*H*) to inhibit serine proteases and cathepsins, respectively. One-way ANOVA was performed for statistical analysis comparing all groups to SARS-PRRA PPs at the highest concentration. * P<0.05. Error bars indicate SEM. PP = pseudoparticles. Representative experiments in triplicate from at least two independent experiments are shown.

### SARS-CoV-2 entry and replication is dependent on serine proteases in differentiated organoid-derived human airway cells

Altogether, our findings show that SARS-CoV-2 preferentially uses serine proteases for entry, when present, and that the multibasic cleavage site increases fusogenicity and infection of human airway cells. Hence, serine protease inhibition could be an attractive therapeutic option. Therefore, we assessed whether camostat could block SARS-CoV-2 entry and replication using differentiated organoid-derived human airway spheroids. In these differentiated spheroids the apical side of the cells was facing outwards (Fig. 5A-B), facilitating virus excretion into the culture medium. These cells were infected with SARS-CoV-2 at a high multiplicity of infection (MOI) of 2, but pretreatment with camostat efficiently blocked virus infection as evidenced by confocal microscopy on spheroids fixed at 16 hours post infection (Fig. 5A). Cathepsin inhibition did not affect entry. At 24 hours post infection, SARS-CoV-2 infection spread in organoids treated with DMSO or E64D, but only rare single cells were observed after camostat treatment (Fig. 5B). Next, we tested whether virus replication was affected by camostat pretreatment of the airway spheroids. After infection at a MOI of 2, replication was assessed at 2, 24, and 48 hours post infection by RT-qPCR and live virus titration. In the control spheroids, SARS-CoV-2 replicated to high titers, while camostat reduced replication by approximately 90% (Fig. 5C-E). We also tested the effect of camostat in 2D differentiated airway cultures at air-liquid interface using a low MOI of 0.1. Here, viral titers in apical washes did not increase after camostat pretreatment (Fig. 5F), whereas replication to moderate titers was observed in the control wells. These findings indicate that SARS-CoV-2 utilizes serine proteases for efficient entry into relevant human airway cells and serine protease inhibition decreases replication.

**Figure 5.**
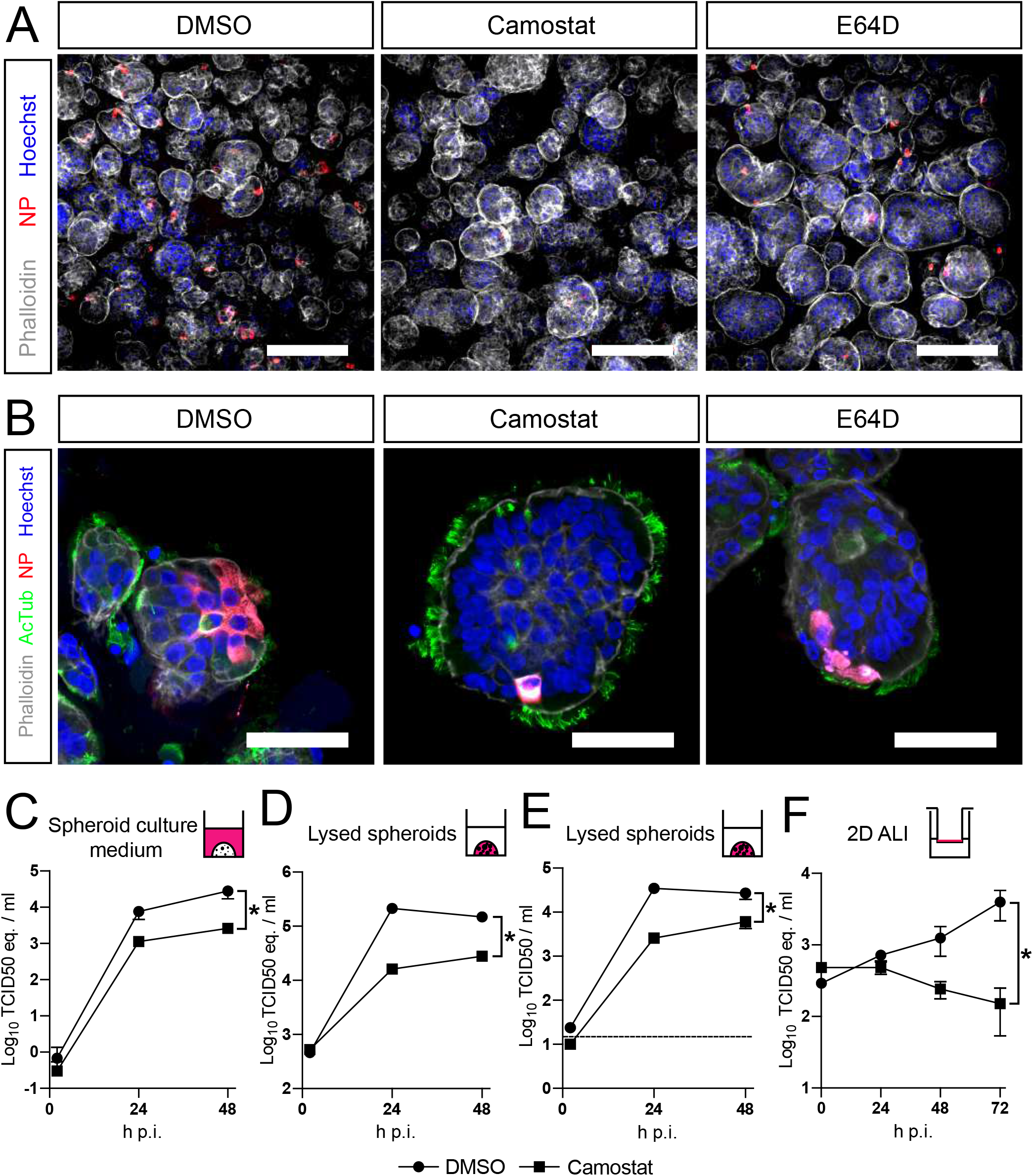
SARS-CoV-2 entry and replication is dependent on serine proteases in differentiated organoid-derived human airway cells. (*A* and *B*) Differentiated bronchiolar (A) or bronchial (B) airway spheroid cultures were infected at a MOI of 2. 16 hours (*A*) or 24 hours (*B*) post infection they were fixed and stained for viral nucleoprotein (red). Nuclei were stained with hoechst (blue) and actin was stained using phalloidin (white). AcTub stains ciliated cells (green). Scale bars indicate 200 μm in *A* and 50 μm in *B*. Representative images are shown from two independent experiments. (*C* to *E*) Replication kinetics of SARS-CoV-2 in bronchiolar airway spheroid cultures pretreated with camostat or carrier (DMSO). (*C* and *D*) TCID50 equivalents (eq.) per ml are shown in culture medium (*C*) and lysed organoids (*D*). (*E*) Live virus titers (TCID50/ml) in lysed organoids. Dotted line indicates limit of detection. (*F*) Replication kinetics of SARS-CoV-2 in 2D tracheal air-liquid interface airway cultures pretreated with camostat or carrier (DMSO). TCID50 eq. per ml in apical washes are shown. Error bars indicate SEM. H p.i. = hours post infection. Two-way ANOVA was performed for statistical analysis. * P<0.05.

## Discussion

SARS-CoV-2 harbors a remarkable multibasic cleavage site in its S protein. Recent findings show that replacing this site with the SARS-CoV monobasic cleavage site decreases PP infectivity on the adenocarcinoma cell line Calu-3, suggesting that this motif is a human airway adaptation (18). This raised the question whether similar findings would be obtained in relevant lung cells. In this study, we found that the SARS-CoV-2 multibasic cleavage site alters tropism by increasing infectivity on differentiated organoid-derived human airway spheroids. Furthermore, we report that the multibasic cleavage site increases S protein fusogenicity, entry rate and serine protease usage. Blocking serine protease activity in organoid-derived differentiated human airway spheroids using the clinically approved drug camostat mesylate effectively inhibited SARS-CoV-2 entry and replication, suggesting that serine protease-mediated entry is the main entry route *in vivo*.

In contrast to SARS-CoV-2, SARS-CoV does not contain a multibasic cleavage site, yet infects Calu-3 cells with similar efficiency. Introducing a multibasic cleavage site to SARS-CoV S did not increase infectivity, indicating that the SARS-CoV S has other adaptations to enter airway cells. These data are in agreement with a study that observed no benefit of furin cleavage on SARS-CoV infectivity (22). Whereas SARS-CoV-2 appears to have adapted to increase fusogenicity and serine protease-mediated S activation for rapid plasma membrane entry, SARS-CoV may have specific adaptations to enter these cells more slowly. Slower viral dissemination may explain why most SARS-CoV patients entered the infectious phase of the disease after symptom onset (23, 24). This could have played a role in the 2003 SARS-CoV epidemic, allowing strict public health interventions including quarantining of symptomatic people and contact tracing to halt viral spread. For SARS-CoV-2, however, several studies have reported that individuals can transmit the virus to others before they become symptomatic (25–29). Whether differences in entry rate allow SARS-CoV-2 to spread more efficiently in the human airway compared with SARS-CoV remains to be investigated. It will be interesting to assess this using authentic SARS-CoV-2 containing multibasic cleavage site mutations, which requires a reverse genetic system, not available to this study at present. Whether cell-cell fusion also plays a role in virus dissemination needs to be determined. In a cell-cell fusion assay and in organoid-derived human airway cells cultured at air-liquid interface we show that SARS-CoV-2 is more fusogenic than SARS-CoV and that fusogenicity is increased by the SARS-CoV-2 S multibasic cleavage site. The role of cell-cell fusion in coronavirus transmission and pathogenesis has not been investigated in detail, but it could be a strategy to avoid extracellular immune surveillance and may increase the viral dissemination rate in the airways *in vivo*. Whether the multibasic cleavage site also affects entry into cells of other organs needs to be investigated further. Although SARS-CoV-2 symptoms are mainly respiratory, recent reports indicate frequent extrapulmonary manifestations, including but not limited to thrombotic complications, acute kidney injury, gastrointestinal symptoms, and dermatologic complications (30). Of note, acute kidney injury was uncommon during the SARS-CoV epidemic (31). It is unclear at this moment whether these manifestations are the result of extrapulmonary viral replication.

Using VeroE6-TMPRSS2 cells that have both active serine protease- and cathepsin-mediated entry pathways, we show that the multibasic cleavage site increases serine protease-mediated S activation, while decreasing cathepsin-mediated S activation. This indicates that the multibasic cleavage site could be an adaptation to serine protease-mediated entry. Whether this site improves S activation by any protease or by serine proteases specifically remains to be tested. Encountering serine proteases first may result in more plasma membrane entry over endosomal entry. More efficient fusion of multibasic motif containing S proteins may be caused by increased S2’ cleavage due to higher accessibility of a S1/S2 cleaved S compared with an uncleaved S. S1/S2 cleavage was recently shown to increase the binding of S to ACE2 (32). Structural changes caused by S1/S2 cleavage may affect protease accessibility as well and may increase subsequent S2’ cleavage.

While mutations in the SARS-CoV-2 multibasic cleavage site decreased airway cell infectivity, they increased infectivity on VeroE6 cells. Several groups have reported mutations or deletions in or around the SARS-CoV-2 multibasic cleavage site that arise in cell culture on VeroE6 cells (33–35), indicating that the lack of a multibasic cleavage site creates a selective advantage in cell culture on VeroE6 cells. The mechanism behind this remains unknown. The increased infectivity of multibasic cleavage site mutants was not observed by Hoffmann and colleagues (2020), but in that study complete cleavage motifs including four amino acids N-terminally from the minimal RXXR cleavage site were exchanged between SARS-CoV-2 and SARS-CoV (18). In contrast, we mutated single sites or removed/inserted only the PRRA motif. Importantly, a cell culture adapted virus containing a complete deletion of the multibasic cleavage site was recently shown to be attenuated in hamsters (35). These studies support our findings that the SARS-CoV-2 multibasic cleavage site affects tropism, facilitates airway cell entry and show that proper characterization of virus stocks is essential.

Entry inhibition has been proposed as an effective treatment option for SARS-CoV-2. Chloroquine can block SARS-CoV and SARS-CoV-2 entry *in vitro* into VeroE6 cells (36, 37), but does not block entry into cells expressing serine proteases (Calu-3 and Vero-TMPRSS2) (38). This is expected, as chloroquine acts in the endosome, while the endosomal entry pathway is not utilized in serine protease expressing cells. As lung cells express serine proteases, inhibitors that block endosomal entry are likely to be ineffective *in vivo*. These findings highlight that drug screens should be performed directly in relevant cells to prevent wasting resources. Our study shows that SARS-CoV-2 replication in human airway spheroids infected with a high MOI is inhibited ~90% by camostat, suggesting that this drug may be effective *in vivo*. Future studies assessing the efficacy and safety of camostat in animal models should be conducted. For SARS-CoV, camostat improved survival to 60% in a lethal mouse model (39). In the same study, inhibition of cathepsins using a cysteine protease inhibitor was ineffective, supporting a critical role for serine proteases in viral spread and pathogenesis *in vivo*. In Japan, camostat has been clinically approved to treat chronic pancreatitis, and thus represents a potential therapy for respiratory coronavirus infections.

Overall, our findings indicate that the multibasic cleavage motif in the SARS-CoV-2 S protein facilitates serine protease-mediated airway cell entry, increasing pandemic potential. In addition, our findings using authentic SARS-CoV-2 in a relevant human airway model suggest that serine protease inhibition is an effective antiviral strategy, as either a therapy or prophylaxis.

## Materials and Methods

### Viruses and cells

Vero, VeroE6, and VeroE6 stable cell lines were maintained in Dulbecco’s modified Eagle’s medium (DMEM, Gibco) supplemented with 10% fetal calf serum (FCS), HEPES (20mM, Lonza), sodium bicarbonate (0.075%, Gibco), penicillin (100 IU/mL) and streptomycin (100 IU/mL) at 37°C in a humidified CO_2_ incubator. Calu-3 and Calu-3 stable cell lines were maintained in Eagle’s minimal essential medium (EMEM, ATCC®) supplemented with 20% FCS, penicillin (100 IU/mL) and streptomycin (100 IU/mL) at 37°C in a humidified CO_2_ incubator. HEK-293T cells were cultured in DMEM supplemented with 10% fetal calf serum (FCS), sodium pyruvate (1mM, Gibco), non-essential amino acids (1X, Lonza), penicillin (100 IU/mL) and streptomycin (100 IU/mL) at 37°C in a humidified CO_2_ incubator. TMPRSS2 and GFP1-10 overexpression cells were maintained in medium containing hygromycin (Invitrogen) and geneticin (Invitrogen), respectively. SARS-CoV-2 (isolate BetaCoV/Munich/BavPat1/2020; European Virus Archive Global #026V-03883; kindly provided by Dr. C. Drosten) and SARS-CoV (isolate HKU39849) were propagated on Vero cells in Opti-MEM I (1X) + GlutaMAX (Gibco), supplemented with penicillin (100 IU/mL) and streptomycin (100 IU/mL) at 37°C in a humidified CO_2_ incubator. The SARS-CoV-2 isolate was obtained from a clinical case in Germany, diagnosed after returning from China. Stocks were produced by infecting cells at a MOI of 0.01 and incubating the cells for 72 hours. The culture supernatant was cleared by centrifugation and stored in aliquots at −80°C. Stock titers were determined by preparing 10-fold serial dilutions in Opti-MEM I (1X) + GlutaMAX. Aliquots of each dilution were added to monolayers of 2 × 10^4^ VeroE6 cells in the same medium in a 96-well plate. Plates were incubated at 37°C 5% CO_2_ for 5 days and then examined for cytopathic effect. The TCID50 was calculated according to the method of Spearman & Kärber. All work with infectious SARS-CoV and SARS-CoV-2 was performed in a Class II Biosafety Cabinet under BSL-3 conditions at Erasmus Medical Center.

### Isolation, culture and differentiation of human airway stem cells

Adult lung tissue was obtained from residual, tumor-free, material obtained at lung resection surgery for lung cancer. The Medical Ethical Committee of the Erasmus MC Rotterdam granted permission for this study (METC 2012-512). Isolation, culture and differentiation was performed as described previously (40) according to a protocol adapted from Sachs and colleagues (2019) (21). Differentiation time on air-liquid interphase was 10-11 weeks. For this study, we used carefully dissected out bronchial material for the generation of bronchial airway organoids. Bronchiolar organoids were generated from distal lung parenchymal material. Tracheal stem cells were collected from tracheal aspirates of intubated preterm infants (<28 weeks gestational age) (41) and cultured as described before (21). For the tracheal aspirates, informed consent was obtained from parents and approval was given by the Medical Ethical Committee (METC no. MEC-2017-302). All donor materials were completely anonymized.

### Authentic virus infection of primary airway cells

To assess differences in syncytium formation, 2D air-liquid interface differentiated airway cultures were washed three times with 500 μL advanced DMEM/F12 (AdDF+++, Gibco) before inoculation from the apical side at a MOI of 1 in 200 μL AdDF+++ per well. Next, cultures were incubated at 37°C 5% CO_2_ for 2 hours before washing 4 times in 500 μL AdDF+++. Cultures were washed daily from the apical side with 300 μL AdDF+++ to facilitate virus spread. At 72 hours post infection, cells were fixed for immunofluorescent staining.

To determine the effect of camostat on SARS-CoV-2 entry, we incubated bronchial or bronchiolar cultures that were differentiated at air-liquid interface for 10-11 weeks with 100% dispase in the basal compartment of a 12 mm Transwell insert. After a 10 minute incubation step at 37°C 5% CO_2_, dispase was removed and cold 500 μL AdDF+++ was pipetted onto the apical side of the Transwell to dislodge the pseudostratified epithelial layer, which was subsequently mechanically sheared by pipetting using a P1000 tip. The resulting epithelial fragments were washed twice in 5 ml AdDF+++ before treatment with 10μM camostat, 10μM E64D or carrier (DMSO) in Pneumacult (PC) ALI medium (Stemcell) on ice for 1 hour. Next, fragments were infected at a MOI of 2 for 2 hours at 37°C 5% CO_2_ in the presence of inhibitors or DMSO. Subsequently, fragments were washed three times in 5 ml cold AdDF+++ before being embedded in 30 μL BME (Type 2; R&D Systems) per well in a 48-well plate. Approximately 200000 cells were plated per well. After solidification of the BME, 200 μL PC was added per well and plates were incubated at 37°C 5% CO_2_.

To assess SARS-CoV-2 replication in the presence of camostat, bronchiolar airway spheroids or 2D air-liquid interface differentiated tracheal airway cultures were infected as described above. For spheroids, culture medium was collected at the indicated time points and frozen at −80°C. After culture medium collection, BME droplets containing spheroids were resuspended in 200 μL AdDF+++ and samples were frozen at −80°C to lyse the cells. To assess 2D air-liquid interface differentiated airway culture replication kinetics, apical washes were collected at the indicated time points by adding 200 μL AdDF+++ apically, incubating for 15 minutes at 37°C 5% CO_2_ and collecting the sample before storage at −80°C. For virus titrations using RT-qPCR (38) or TCID50 determination, samples were thawed and centrifuged at 500 x g for 3 min. For TCID50 determination, six replicates were performed per sample.

### Fixed immunofluorescence microscopy and immunohistochemistry

Transwell inserts were fixed in formalin, permeabilized in 70% ethanol, and blocked for 60 minutes in 10% normal goat serum or 3% bovine serum albumin (BSA) in PBS (blocking buffer). For organoids 0.1% triton X-100 was added to the blocking buffer to increase antibody penetration. Cells were incubated with primary antibodies overnight at 4°C in blocking buffer, washed twice with PBS, incubated with corresponding secondary antibodies Alexa488-, 594 and 647-conjugated secondary antibodies (1:400; Invitrogen) in blocking buffer for 2 hours at room temperature, washed two times with PBS, incubated with indicated additional stains (TO-PRO3, phalloidin-633 (SC-363796, Santa Cruz Biotechnology), Hoechst), washed twice with PBS, and mounted in Prolong Antifade (Invitrogen) mounting medium.

SARS-CoV-2 and SARS-CoV were stained with mouse-anti-SARS-CoV nucleoprotein (40143-MM05, 1:400, Sino Biological) or rabbit-anti-SARS-CoV nucleoprotein (40143-T62, 1:400, Sino biological). Tight junctions were stained using mouse-anti-ZO1 (ZO1-1A12, 1:200, Invitrogen). Club cells and goblet cells were stained with mouse-anti-CC10 (sc-390313 AF594, 1:100, Santa Cruz Biotechnology) and mouse anti-MUC5AC (MA5-12178, 1:100, Invitrogen), respectively. Ciliated cells were stained with mouse-anti-FOXJ1 (14-9965-82, 1:200, eBioscience) and mouse-anti-AcTub (sc-23950 AF488, 1:100, Santa Cruz Biotechnology). For lineage marker stainings formalin-fixed inserts were paraffin-embedded, sectioned and deparaffinized as described before prior to staining (42). Samples were imaged on a LSM700 confocal microscope using ZEN software (Zeiss). Representative images were acquired and shown as Z-projections, single slices or XZ cross sections.

Immunohistochemistry was performed as described previously (42) on formalin fixed, paraffin embedded Transwell inserts. ACE2 and TMPRSS2 were stained using goat-anti-hACE2 (AF933, 1:200, R&D Systems) and mouse-anti-TMPRSS2 (sc-515727, 1:200, Santa Cruz Biotechnology), and visualized with rabbit-anti-goat (P0160, 1:200, Dako) and goat-anti-mouse (PO260, 1:100, Dako) horseradish peroxidase labeled secondary antibody, respectively. Samples were counterstained using haematoxylin.

### GFP-complementation fusion assay

HEK-293T cells were grown in 6-well format to 70-80% confluency and were transfected with 1.5 μg pGAGGS-spike (all coronavirus S variants described above) DNA and pGAGGS-β-Actin-P2A-7xGFP11-BFP DNA or empty vector DNA with PEI in a ratio of 1:3 (DNA:PEI). Beta-actin was tagged with 7xGFP11 expressed in tandem and blue fluorescent protein (BFP). The two genes were separated by a P2A self-cleaving peptide. Two variants of this construct were used. One variant contained a GSG linker located N-terminally from the P2A site to improve self-cleavage and this construct was used in qualitative confocal microscopy experiments. A variant lacking the GSG linker was less efficiently cleaved as indicated by both cytoplasmic and nuclear localized GFP, but this generated an equal distribution of GFP throughout the cell and therefore it was used for all fusion assays in which the sum of all GFP+ pixels was calculated. Transfected HEK-293T cells were incubated overnight at 37°C 5% CO_2_. GFP1-10 expressing cells were seeded in a 12-well plate to achieve 90-100% confluency after overnight incubation at 37°C 5% CO_2_ and medium was refreshed with Opti-MEM I (1X) + GlutaMAX. HEK-293T cells were resuspended in PBS by pipetting to generate a single cell suspension and added to GFP1-10 expressing cells in a ratio of 1:80 (HEK-293T cells: GFP1-10 expressing cells). Fusion events were quantified by detecting GFP+ pixels after 18 hours incubation at 37°C 5% CO_2_ using Amersham™ Typhoon™ Biomolecular Imager (channel Cy2; resolution 10μm; GE Healthcare). Data was analyzed using the ImageQuant TL 8.2 image analysis software (GE Healthcare) by calculating the sum of all GFP+ pixels per well. For nuclear counting fluorescence microscopy images were obtained with a Carl ZEISS Vert.A1 microscope paired with an AxioCam ICm1 camera and Colibri 7 laser (469/38nm for GFP and 365/10nm for BFP) using ZEN analysis software (20x magnification). Nuclei per syncytia were calculated by counting BFP-positive nuclei after 18 hours incubation at 37°C 5% CO_2_. Confocal microscopy images were taken on a LSM700 confocal microscope using ZEN software. Representative images were acquired and shown as single slices.

### Coronavirus S pseudotyped particle production

For the production of SARS-CoV and SARS-CoV-2 S PPs, as well as multibasic cleavage site mutant PPs, HEK-293T cells were transfected with 15 μg S expression plasmids. Twenty-four hours post-transfection, the medium was replaced for Opti-MEM I (1X) + GlutaMAX, and cells were infected at a MOI of 1 with VSV-G PPs. Two hours post-infection, cells were washed three times with Opti-MEM I (1X) + GlutaMAX and replaced with medium containing anti-VSV-G neutralizing antibody (clone 8G5F11; Absolute Antibody) at a dilution of 1:50000 to block remaining VSV-G PPs. The supernatant was collected after 24 hours, cleared by centrifugation at 2000 x g for 5 minutes and stored at 4°C until use within 7 days. Coronavirus S PPs were titrated on VeroE6 cells as described in the supplementary information (SI) appendix.

### Entry route assay

VeroE6, Calu-3 and VeroE6-TMPRSS2 cells were seeded in 24 well plates and kept at 37°C 5% CO_2_ overnight to achieve 80-100% confluency by the next day. Cells were pretreated with a concentration range of camostat, E64D or DMSO (with all conditions containing equal concentrations of DMSO) in Opti-MEM I (1X) + GlutaMAX for 2 hours before infecting with on average 1000 PPs per well. Plates were incubated overnight at 37°C 5% CO_2_ before scanning for GFP signal as described above.

### Entry speed assay

Calu-3 cells were seeded as for entry route assays and pre-treated with 10μM E64D. After one hour, PPs were added per well to achieve 1000 infected cells in the control well. At the same time as addition of PPs, 10μM camostat was added into the first set of wells (t=0). DMSO was added to controls. The same inhibitor was added in the next sets of wells in triplicate 2, 4 and 6 hours post infection. Plates were incubated overnight at 37°C 5% CO_2_ before scanning for GFP signal as described above.

Authentic virus entry speed was performed in the same manner, by infecting Calu-3 cells with 1×10^4^ TCID50 SARS-CoV-2 and 5×10^4^ TCID50 SARS-CoV. After 12 hours, plates were fixed and blocked as above for transwell inserts. Cells were incubated with mouse-anti-double stranded RNA (Clone J2, 1:500, Scicons) in blocking buffer for 2 hours at room temperature or overnight at 4°C. Cells were washed twice with PBS and stained with Alexa488 conjugated secondary antibody (1:500 Invitrogen) in blocking buffer for an hour at room temperature. Finally, cells were washed twice with PBS and scanned in PBS on the Amersham™ Typhoon as described above.

### Coronavirus S pseudotyped particle concentration

PPs were concentrated on a 10% sucrose cushion (10% sucrose, 15mM Tris-HCl, 100mM NaCl, 0.5mM EDTA) at 20000g for 1.5 hours at 4°C. Supernatant was decanted and pellet was resuspended overnight at 4°C in Opti-MEM I (1X) + GlutaMAX to achieve 100-fold concentration. PPs were titrated and aliquots were lysed in 1X Laemmli buffer (Bio-Rad) containing 5% 2-mercaptoethanol for western blot analysis.

### Pseudoparticle infection of primary airway cells

To determine the effect of multibasic cleavage site mutations on SARS-CoV-2 entry, we obtained airway culture fragments from 2D differentiated bronchiolar cultures as described above. Next, fragments were infected with equal volumes of concentrated wild type and multibasic cleavage site mutant PPs for 2 hours at 37°C 5% CO_2_. Subsequently, the supernatant was replaced with 30 μL BME and plated in a 48-well plate. Approximately 200000 cells were plated per well. After solidification of the BME, 200 μL PC was added per well and plates were incubated at 37°C 5% CO_2_. After overnight incubation, the amount of infected cells and organoids per field were counted and images taken using a Carl ZEISS Vert.A1 microscope paired with an AxioCam ICm1 camera and Colibri 7 laser (469/38nm for GFP) using ZEN analysis software.

### Statistical analysis

Statistical analysis was performed with the GraphPad Prism 5 and 8 software using a t-test, one way ANOVA or two-way ANOVA followed by a Bonferroni multiple-comparison test.

Additional experimental methods, including cloning, stable cell line generation, VSV delta G rescue, western blotting and silver staining, can be found in the SI appendix.

## Acknowledgments

This work was supported by NWO Grant 022.005.032, partly financed by the Netherlands Organization for Health Research and Development (ZONMW) grant agreement 10150062010008 to B.L.H and co-funded by the PPP Allowance (grant agreement LSHM19136) made available by Health Holland, Top Sector Life Sciences & Health, to stimulate public-private partnerships.

## Notes

### Competing Interest Statement

The authors have declared no competing interest.

