## Supplementary Material for "The SARS-CoV-2 multibasic cleavage site facilitates early serine protease-mediated entry into organoid-derived human airway cells"

**This PDF file includes:**

Supplementary methods
Figure legends S1 to S3
Supplementary references
Figures S1 to S3

**Supplementary methods**

*Cloning*

Codon-optimized SARS-CoV (isolate CUHK-W1; VG40150-G-N) S expression plasmids (pCMV) were ordered from Sino-Biological and subcloned into pCAGGS using the ClaI and KpnI sites. The last 19 amino acids containing the golgi retention signal of the SARS-CoV S protein were deleted to enhance PP production. Codon-optimized cDNA encoding SARS-CoV-2 S glycoprotein (isolate Wuhan-Hu-1) with a C-terminal 19 amino acid deletion was synthesized and cloned into pCAGSS in between the EcoRI and BglII sites. S expressing pCAGGS vectors were used for GFP-complementation fusion assays and equivalent S proteins with the C-terminal deletion were used for the production of PPs, as described in the material and methods. The cDNA encoding GFP1-10 was obtained from Addgene and was subcloned into pQXCIN (Clontech) in between BamHI and EcoRI to obtain the pQXCIN-GFP1-10 vector. The cDNA encoding human TMPRSS2 (NM_005656; OHu13675D) was obtained from Genscript. The cDNA fused to a C-terminal HA tag was subcloned into pQXCIH (Clontech) in between the NotI and PacI sites to obtain the pQXCIH-TMPRRS2-HA vector. A synthetic DNA construct of β-Actin-7xGFP11-P2A-BFP was ordered from GenScript and subcloned into pGAGGS using EcoRI and BglII. SARS-CoV and SARS-CoV-2 S protein mutations (SARS-PRRA, SARS-2-Del-PRRA, SARS-2-R685A, SARS-2-R685H) were generated by subcloning synthetic DNA constructs (Genscript) containing the desired mutations into the pCAGGS-S vectors or by mutagenesis PCR.

*Generation of stable cell lines expressing GFP1-10 and TMPRSS2*

VeroE6 GFP1-10, VeroE6-TMPRSS2 cells, VeroE6-TMPRSS2 GFP1-10 cells and Calu-3 GFP1-10 cells were generated by retroviral transduction. To produce the retrovirus, 10 µg pQXCIH-TMPRRS2-HA or pQXCIN-GFP1-10 was co-transfected with polyethylenimine (PEI) with 6.5 µg pBS-gag-pol (Addgene #35614) and 5 µg pMD2.G (Addgene #12259) in a 10 cm dish of 70% confluent HEK-293T cells in Opti-MEM I (1X) + GlutaMAX. Retroviral particles were harvested at 72 hours post transfection, cleared by centrifugation at 2000 x g, filtered through a 0.45µm low protein binding filter (Millipore), and used to transduce designated cells. Polybrene (Sigma) was added at a concentration of 4 µg/ml to enhance transduction efficiency. Transduced cells were selected with hygromycin B (Invitrogen) for TMPRSS2 cells and/or geneticin (Invitrogen) for GFP1-10 cells.

*VSV delta G rescue*

The protocol for VSV-G PP rescue was adapted from Whelan and colleagues (1995) (1). VSV rescue plasmids pVSV-eGFP-dG (#31842), pMD2.G (#12259), pCAG-VSV-P (#64088), pCAG-VSV-L (#64085), pCAG-VSV-N (#64087) and pCAGGS-T7Opt (#65974) were ordered from Addgene. Briefly, a 70% confluent 10 cm dish of HEK-293T cells was transfected with 10µg pVSV-eGFP-dG, 2µg pCAG-VSV-N (nucleocapsid), 2µg pCAG-VSV-L (polymerase), 2µg pMD2.G (glycoprotein, VSV-G), 2µg pCAG-VSV-P (phosphoprotein) and 2µg pCAGGS-T7Opt (T7 RNA polymerase) using PEI at a ratio of 1:3 (DNA:PEI) in Opti-MEM I (1X) + GlutaMAX. Forty-eight hours post transfection, the supernatant was transferred onto new plates transfected 24 hours prior with VSV-G. After a further 48 hours, these plates were re-transfected with VSV-G. After 24 hours the resulting PPs were collected, cleared by centrifugation at 2000 x g for 5 minutes, and stored at -80°C. Subsequent VSV-G PP batches were produced by infecting VSV-G transfected HEK-293T cells with VSV-G PPs at a MOI of 0.1. Titers were determined by preparing 10-fold serial dilutions in Opti-MEM I (1X) + GlutaMAX. Aliquots of each dilution were added to monolayers of $2 \times 10^4$ Vero cells in the same medium in a 96-well plate. Three replicates were performed per PP stock. Plates were incubated at 37°C overnight and then scanned using an Amersham™ Typhoon scanner. Individual infected cells were quantified using ImageQuant TL software. All PP work was performed in a Class II Biosafety Cabinet under BSL-2 conditions at Erasmus Medical Center.

*Spike protein western blot*

Concentrated PPs diluted in 4x Laemmli loading buffer were boiled for 30 minutes at 95°C. S transfected HEK-293T cells were lysed using IP Lysis Buffer (Pierce). Cell lysate was rotated for 30 minutes and centrifuged for 10 minutes at 15000 x g. Supernatant was used for subsequent protein expression analysis. Lysates were diluted in 4x Laemmli loading buffer containing 20% 2-mercaptoethanol and boiled for 30 minutes at 95°C. PPs and cell lysates were used for SDS-PAGE analysis using precast 10% TGX gels (Bio-Rad). Gels were run in tris-glycine SDS (TGS) buffer at 50V for 30 minutes and subsequently at

120V for 90 minutes. Transfer was performed at 300mA for 55 minutes onto .45 µm Immobilon-FL PVDF Membranes in tris-glycine buffer containing 20% methanol. Spike was stained using polyclonal rabbit-anti-SARS-CoV S1 (1:1000, Sino Biological) followed by infrared labelled secondary antibodies (1:20000, Licor). All cell lysate western blots are stained for GAPDH using a monoclonal mouse-anti-GAPDH antibody (sc-32233, 1:1000, Santa Cruz Biotechnology) followed by infrared labelled secondary antibodies. Western blots were scanned on an Odyssey CLx and analyzed using Image Studio Lite Ver 5.2 software.

*Silver staining*

All PP western blots had corresponding silver stains performed to assess the quality of the PP preps and for the detection of VSV-N. Samples boiled in Laemmli buffer for western blot analysis were also ran on a 10% w/v gel at 50V for 30 minutes followed by 120V for 90 minutes before transferring gel into ultrapure water. Silver stains were performed per manufacturer's instructions using the Silver Stain for Mass Spectrometry kit (Pierce). Colorimetric images were taken on the Amersham™ AI600 (GE Healthcare).

**Fig. S1. Organoid-derived 2D air-liquid interface cultures are well-differentiated and express ACE2 and TMPRSS2.** (*A* to *C*) Immunofluorescent or immunohistochemistry staining of differentiated airway cultures. Anti-AcTub (green) and anti-FOXJ1 (white) stains ciliated cells (*A*), anti-SCGB1A1 (magenta) stains club cells (*B*) and anti-MUC5AC (yellow) stains goblet cells (*C*). Nuclei are stained with hoechst (blue). (*D* to *E*) Airway cultures also expressed the SARS-CoV-2 entry receptor ACE2 (*D*) and TMPRSS2 (*E*). Haematoxylin was used as a counterstain in *D* and *E*. Scale bars indicate 20 µm. Representative images are shown from a bronchiolar culture.

**Fig. S2. SARS-CoV PP infectivity into Calu-3 cells is not altered by the insertion of the multibasic cleavage site.** Titrations of SARS-CoV PPs and SARS-PRRA PPs on Calu-3 cells. Error bars indicate SEM. A representative experiment in triplicate from three independent experiments is shown.

**Fig. S3. A GFP-complementation based assay for assessing coronavirus fusogenicity.** (*A*) HEK-293T cells expressing an empty vector or S protein together with GFP-11 tagged beta actin and a BFP containing a nuclear localization signal were added to cells stably expressing GFP1-10. Fusion of these two cell types allowed GFP-complementation in cells expressing a nuclear BFP, facilitating easy quantification of nuclei per syncytial cell. Unfused cells only expressed BFP in the nucleus. Fusion with VeroE6 GFP1-10 cells 18 hours after addition of the fusogenic HEK-293T is shown as an example. (*B* to *D*) Full well scans of the complemented GFP signal 18 hours after addition of the fusogenic HEK-293T cells to Calu-3 GFP1-10 (*B*), VeroE6 GFP1-10 (*C*) and VeroE6-TMPRSS2 GFP1-10 (*D*) cells are shown. Dashed areas are enlarged next to each well. Scale bars indicate 50 μm.

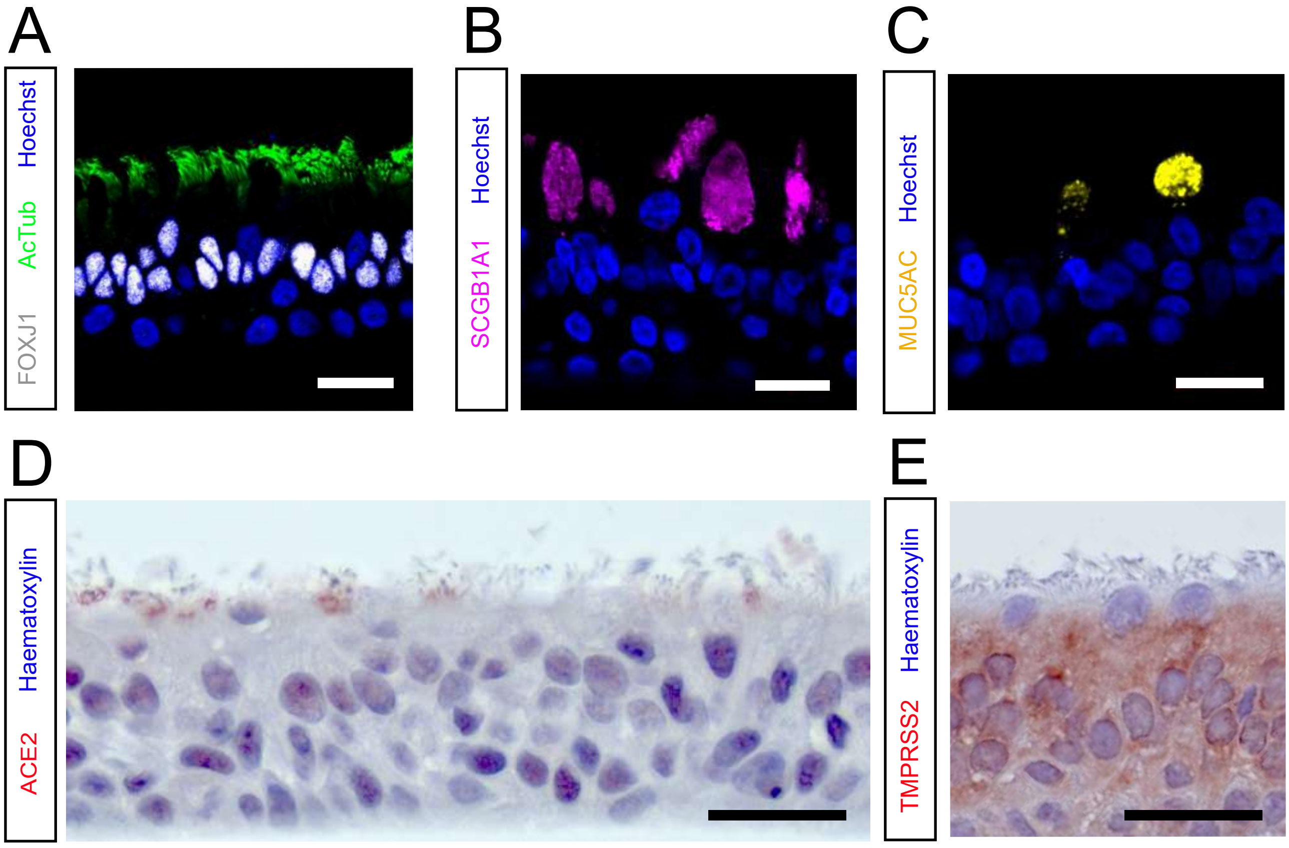


**Fig. S1. Organoid-derived 2D air-liquid interface cultures are well-differentiated and express ACE2 and TMPRSS2.** (*A* to *C*) Immunofluorescent or immunohistochemistry staining of differentiated airway cultures. Anti-AcTub (green) and anti-FOXJ1 (white) stains ciliated cells (*A*), anti-SCGB1A1 (magenta) stains club cells (*B*) and anti-MUC5AC (yellow) stains goblet cells (*C*). Nuclei are stained with hoechst (blue). (*D* to *E*) Airway cultures also expressed the SARS-CoV-2 entry receptor ACE2 (*D*) and TMPRSS2 (*E*). Haematoxylin was used as a counterstain in *D* and *E*. Scale bars indicate 20 μm. Representative images are shown from a bronchiolar culture.

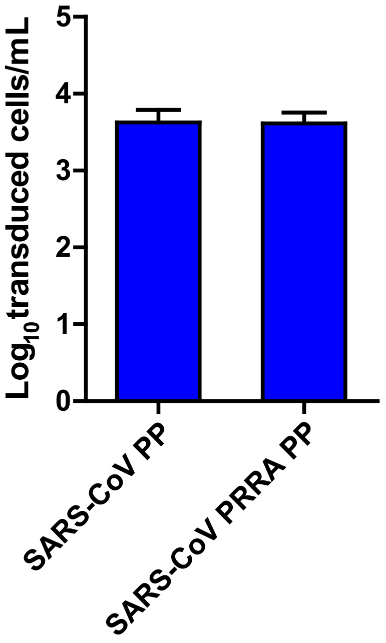


**Fig. S2. SARS-CoV PP infectivity into Calu-3 cells is not altered by the insertion of the multibasic cleavage site.** Titrations of SARS-CoV PPs and SARS-PRRA PPs on Calu-3 cells. Error bars indicate SEM. A representative experiment in triplicate from three independent experiments is shown.

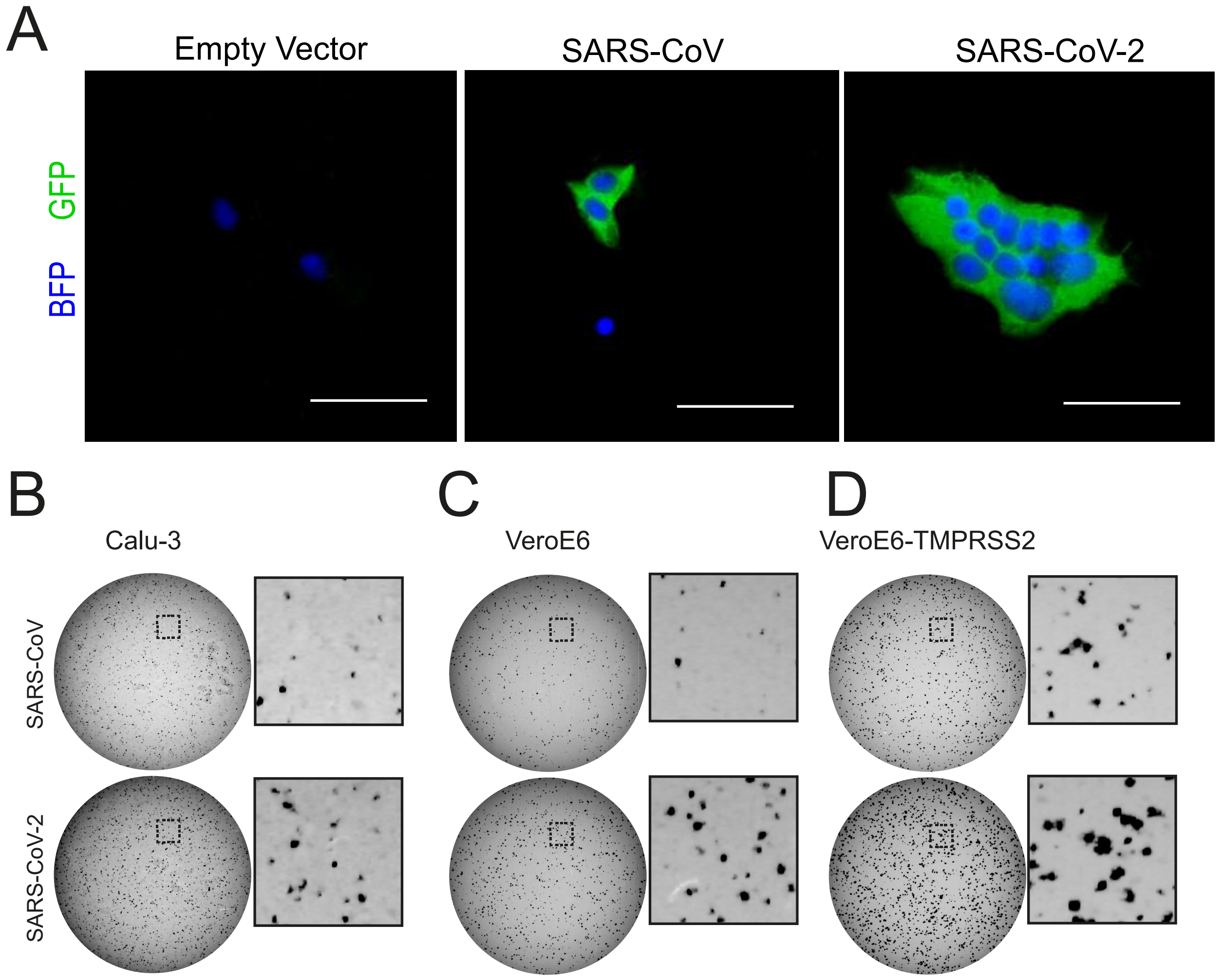


**Fig. S3. A GFP-complementation based assay for assessing coronavirus fusogenicity.** (*A*) HEK-293T cells expressing an empty vector or S protein together with GFP-11 tagged beta actin and a BFP containing a nuclear localization signal were added to cells stably expressing GFP1-10. Fusion of these two cell types allowed GFP-complementation in cells expressing a nuclear BFP, facilitating easy quantification of nuclei per syncytial cell. Unfused cells only expressed BFP in the nucleus. Fusion with VeroE6 GFP1-10 cells 18 hours after addition of the fusogenic HEK-293T is shown as an example. (*B* to *D*) Full well scans of the complemented GFP signal 18 hours after addition of the fusogenic HEK-293T cells to Calu-3 GFP1-10 (*B*), VeroE6 GFP1-10 (*C*) and VeroE6-TMPRSS2 GFP1-10 (*D*) cells are shown. Dashed areas are enlarged next to each well. Scale bars indicate 50 µm.
